# HIF-α signaling regulates the macrophage inflammatory response during *Leishmania major* infection

**DOI:** 10.1101/2024.08.27.605844

**Authors:** Lucy G. Fry, Charity L. Washam, Hayden Roys, Anne K. Bowlin, Gopinath Venugopal, Jordan T. Bird, Stephanie D. Byrum, Tiffany Weinkopff

**Author notes:** Corresponding author: Tiffany Weinkopff Department of Microbiology and Immunology College of Medicine University of Arkansas for Medical Sciences 4301 W. Markham St., Mail Slot 511 Little Rock, AR 72205.

## Abstract

Cutaneous leishmaniasis (CL) contributes significantly to the global burden of neglected tropical diseases, with 12 million people currently infected with *Leishmania* parasites. CL encompasses a range of disease manifestations, from self-healing skin lesions to permanent disfigurations. Currently there is no vaccine available, and many patients are refractory to treatment, emphasizing the need for new therapeutic targets. Previous work demonstrated macrophage HIF-α-mediated lymphangiogenesis is necessary to achieve efficient wound resolution during murine *L. major* infection. Here, we investigate the role of macrophage HIF-α signaling independent of lymphangiogenesis. We sought to determine the relative contributions of the parasite and the host-mediated inflammation in the lesional microenvironment to myeloid HIF-α signaling. Because HIF-α activation can be detected in infected and bystander macrophages in leishmanial lesions, we hypothesize it is the host’s inflammatory response and microenvironment, rather than the parasite, that triggers HIF-α activation. To address this, macrophages from mice with intact HIF-α signaling (LysM^Cre^ARNT^f/+^) or mice with deleted HIF-α signaling (LysM^Cre^ARNT^f/f^) were subjected to RNASequencing after *L. major* infection and under pro-inflammatory stimulus. We report that *L. major* infection alone is enough to induce some minor HIF-α-dependent transcriptomic changes, while infection with *L. major* in combincation with pro-inflammatory stimuli induces numerous transcriptomic changes that are both dependent and independent of HIF-α signaling. Additionally, by coupling transcriptomic analysis with several pathway analyses, we found HIF-α suppresses pathways involved in protein translation during *L. major* infection in a pro-inflammatory environment. Together these findings show *L. major* induces a HIF-α-dependent transcriptomic program, but HIF-α only suppresses protein translation in a pro-inflammatory environment. Thus, this work indicates the host inflammatory response, rather than the parasite, largely contributes to myeloid HIF-α signaling during *Leishmania* infection.

## Introduction

Leishmaniasis is the family of diseases caused by infection with protozoan *Leishmania* parasites. Because pathology depends upon both the species of parasite and the host immune response, leishmaniasis can manifest in three main forms: cutaneous leishmaniasis (CL), mucocutaneous leishmaniasis (MCL), and visceral leishmaniasis (VL). *Leishmania* parasites are transmitted via sandfly bites and are endemic in more than 90 countries across Africa, Asia, and Latin America resulting in 1-2 million new cases of leishmaniasis each year (1,2). There is currently no human vaccine and existing treatments against *Leishmania* parasites are toxic to the host, difficult to administer, require a long duration, and are often ineffective (3). The lack of new treatments or vaccines has made global disease control and elimination efforts challenging, reiterating the importance of understanding the host immune response to identify potential therapeutic targets (4).

Upon a sandfly bite, parasites are taken up by macrophages in the skin. After phagocytosis, parasites reside in phagolysosomes in macrophages and begin multiplying. Controlling parasite burden is dependent on a predominant Th1 immune response where CD4^+^ T cells produce IFNγ which activates macrophages to kill parasites by releasing nitric oxide (NO) and reactive oxygen species (ROS) (5). During CL, neutrophils and inflammatory monocytes are initially recruited to the site of infection (6). Severity of disease is highly dependent upon both parasite burden and the host inflammatory response with excessive inflammation contributing to overall pathology and extending the duration of disease (7). Despite an effective immune response, parasites can persist at low levels in the skin for years even after dermal lesions have resolved (8). Leishmanial lesions are characterized by hypoxia and the presence of pro-inflammatory cells and cytokines (9–12). During inflammatory hypoxia, transcription factors hypoxia-inducible factor (HIF)-1α and HIF-2α are induced by decreased oxygen availability in tissues (13). HIF-α transcription factors are master regulators of genes involved in metabolism and the cellular response to oxygen deprivation (14,15). Upon activation, HIF-α subunits bind aryl hydrocarbon receptor nuclear translocator (ARNT; also known as HIF-1β), and ARNT/HIF-α heterodimers translocate to the nucleus where they induce the transcription of HIF-α target genes (16). HIF-α subunits can also be activated by oxygen-independent mechanisms such as TLR ligation, pro-inflammatory cytokines, or ROS stimulation (17,18). Furthermore, under normoxic conditions LPS induces HIF-1α expression via MyD88/NFκB signaling in macrophages, and mice deficient in HIF-1α are more susceptible to a variety of bacterial and fungal infections (17,19–22).

During CL, human lesions contain elevated levels of HIF-1α and the HIF-α target, VEGF-A (19,23,24). Similarly, HIF-1α and VEGF-A are also elevated in lesions following experimental murine *L. major* infection (25,26). Both inflammatory signaling, such as IFNγ production, as well as hypoxia in the skin promote HIF-1α accumulation in *L. major*-infected macrophages, but which signal occurs first and the relative contributions of each signal to HIF-1α signaling are not known (19,27,28). Myeloid-specific HIF-1α^-/-^ mice infected with *L. major* exhibit increased lesion sizes and parasite burdens due to impaired expression of NOS2, a HIF-1α-specific target gene (19). These data suggest activated HIF-1α in dermal myeloid cells contributes to parasite control through NO production. Additionally, mice deficient in myeloid ARNT/HIF-α signaling (LysM^Cre^ARNT^f/f^; missing both HIF-1α and HIF-2α pathways) infected with *L. major* exhibit decreased myeloid-derived NOS2 and VEGF-A which impairs lymphangiogenesis at the site of infection, resulting in larger lesion sizes, despite parasites being controlled (26). Altogether, these data suggest myeloid HIF-α signaling plays critical roles in both parasite control and lesion resolution during *L. major* infection.

HIF-α activation depends on the *Leishmania* parasite species. In contrast to *L. amazonensis* and *L. donovani* parasites, *L. major* parasites alone do not increase HIF-1α expression or activation under normoxic conditions (11,19,27,29). Additionally, during in vivo *L. major* infection, both infected and bystander macrophages exhibit HIF-α activation compared to macrophages from naïve skin (28). Based on these findings, we hypothesize that during *L. major* infection, it is the host’s inflammatory response and microenvironment, rather than the parasite itself, that triggers HIF-α activation. To address this hypothesis, we performed transcriptomic analyses on macrophages from LysM^Cre^ARNT^f/+^ or LysM^Cre^ARNT^f/f^ that either exhibit intact or impaired ARNT/HIF-α signaling in myeloid cells, respectively. LysM^Cre^ARNT^f/+^ or LysM^Cre^ARNT^f/f^ macrophages were infected or not with *L. major* parasites and then treated or not with LPS and IFNγ to define the importance of ARNT/HIF-α signaling in response to *L. major* parasites in the presence or absence of a pro-inflammatory milieu. We find infection with *L. major* parasites induces transcriptional changes in macrophages and some of these early transcriptomic changes are absent in macrophages without HIF-α signaling. This indicates *L. major* induces some transcriptomic changes that are HIF-α-dependent, and *L. major* infection is sufficient to induce HIF-α activation in vitro, albeit minimal compared to pro-inflammatory stimuli. We discovered under inflammatory conditions, HIF-α signaling suppresses transcripts and pathways involved in translation such as ribosomal transcripts, EIF2 signaling and the ribosome pathway during infection with *L. major.* Additionally, we identified top enriched pathways associated with *L. major* infection during and apart from inflammatory conditions as well as with and without intact HIF-α signaling.

## Materials and Methods

### Parasites

*Leishmania major* strain (WHO/MHOM/IL/80/Friedlin) parasites were cultured with Schneider’s insect media (Gibco) supplemented with 20% heat-inactivated fetal bovine serum (FBS) (Invitrogen), 100 U/mL penicillin/streptomycin (Sigma), and 2mM L-glutamine (Sigma). Metacyclic promastigotes were isolated from 4-5 day old cultures using Ficoll (Sigma) gradient separation for infections (23).

### Mice

C57BL/6 mice were purchased from the National Cancer Institute. Mice with a myeloid-specific *ARNT* conditional knockout were developed by crossing a strain expressing the LysM^Cre^ allele with another strain with a floxed *ARNT* conditional allele and were bred on campus in the vivarium. The LysM^Cre^ARNT^f/f^ and LysM^Cre^ARNT^f/+^ mice were a gift from M. Celeste Simon (University of Pennsylvania, Philadelphia, PA). LysM^Cre^ARNT^f/+^ mice were used as controls for LysM^Cre^ARNT^f/f^ mice. All animals were housed in the vivarium under pathogen-free conditions at the University of Arkansas for Medical Sciences (UAMS). All mice were infected between 6-8 weeks of age and all procedures were approved by UAMS IACUC and followed institutional guidelines.

### Murine Infection in vivo

For dermal ear infections in C57BL/6 mice, 2×10^6^ promastigote *Leishmania major* (WHO/MHOM/IL/80/Friedlin) parasites in 10 µL PBS (Gibco) were injected intradermally into the ear. For analyses, ears were excised, dorsal and ventral sheet were separated. Ear sheets were enzymatically digested for 90 min at 37°C using 0.25 mg/mL Liberase (Roche) and 10 mg/mL DNase I (Sigma) in incomplete RPMI 1640 (Gibco). After digestion, ears were smashed through a filter to obtain a single-cell suspension (28).

### Single-cell RNASequencing Sample Preparation

The scRNASeq samples were prepared and data was acquired as a part of a previous study (28). In short, the Arkansas Children’s Research Institute (ACRI) Genomics and Bioinformatics Core prepared NGS libraries from fresh single-cell suspensions using the 10X Genomics NextGEM 3’ assay for sequencing on the NextSeq 500 platform using Illumina SBS reagents. Trypan Blue exclusion determined cell quantity and viability. Library quality was evaluated with the Advanced Analytical Fragment Analyzer (Agilent) and Qubit (Life Technologies) instruments.

### scRNASeq Data Analysis

Data analysis was performed as a part of a previous study (28). Briefly, the UAMS Genomics Core generated Demultiplexed fastq files which were analyzed using 10X Genomics Cell Ranger alignment and gene counting software, a self-contained scRNASeq pipeline developed by 10X Genomics. The reads were aligned to the mm10 reference transcriptomes using STAR and transcript counts were generated (30,31). The *Seurat* R package processed the raw counts generated by *cellranger count* (32,33). Potential doublets, low quality cells, and cells with a high percentage of mitochondrial genes were filtered out. Cells that have unique feature counts > 75^th^ percentile plus 1.5 times the interquartile range (IQR) or < 25^th^ percentile minus 1.5 time the IQR were filtered. Similarly, cells with mitochondrial counts falling outside the same range for mitochondrial gene percentage were filtered. After filtering, all 8 sequencing runs were merged. The counts were normalized using the LogNormalize method which log-transforms the results(28). Subsequently, the 2000 highest variable features were selected. The data was scaled, and Principal component analysis (PCA) was performed. A JackStraw procedure was implemented to determine the significant PCA components that have a strong enrichment of low p-value features.

A graph-based clustering strategy embedded cells in graph structure (34) Seurat visualized the results in t-distributed stochastic neighbor embedding (tSNE) and Uniform Manifold Approximation and Projection (UMAP) plots (35). Seurat *FindNeighbors* and *FindClusters* functions were optimized to label clusters. Seurat *FindAllMarkers* function finds markers that identify clusters by differential expression, defining positive markers of a single cluster compared to all other cells and comparing those to known markers of expected cell types from previous single-cell transcriptome studies. Cell type determinations were determined by manually reviewing these results, and some clusters were combined if their expression was found to be similar. From here for this work, we specifically provide Feature maps showing transcript expression of HIF-1α, HIF-2α, and corresponding target genes of these transcription factors amongst all clusters, and particularly in macrophages.

### Generation of Bone Marrow-Derived Macrophages

Femurs collected from mice were soaked in 70% ethanol for 2 minutes and then flushed with 10 mL of cDMEM to extract bone marrow cells. Bone marrow cells were counted before plating 5×10^6^ cells per 100 mm Petri dish in 10 mL of conditioned macrophage media (cDMEM with 25% L929 cell supernatants). Cells were cultured for 7 days, refreshing media at day 3. To remove the macrophages from the Petri dish, macrophages were washed with ice-cold PBS and gently removed with a cell scraper. The collected macrophages were counted and loaded into 24-well plates with 1×10^6^ cells in 1 mL cDMEM per well.

### In vitro Infection of BMDM and RNASeq

Bone marrow-derived macrophages (BMDMs) were plated into 24-well plates and allowed to rest overnight. Parasites were added to the wells at a 5:1 multiplicity of infection (MOI). Extracellular parasites were washed away at 2 hours post-infection. After washing, BMDMs were cultured in media with or without 100 ng/mL LPS (Sigma) and 10 ng/mL IFNγ (Peprotech). After 8 hours, the cells washed with PBS, lysed with RLT lysis buffer for RNA extraction, and stored at -80 °C. For transcriptomic RNASeq studies, RNA was extracted following the Qiagen RNEasy Mini-Kit instructions before being subjected to RNASeq analysis. Each experiment group contained 2 or 3 samples for RNASeq analysis.

### RNASeq Analysis

Following demultiplexing, RNA reads were checked for sequencing quality using FastQC (http://www.bioinformatics.babraham.ac.uk/projects/fastqc) and MultiQC (36)(version 1.6). The raw reads were then processed according to Lexogen’s QuantSeq data analysis pipeline with slight modification. Briefly, residual 3’ adapters, polyA read through sequences, and low quality (Q < 20) bases were trimmed using BBTools BBDuk (version 38.52) (https://sourceforge.net/projects/bbmap/). The first 12 bases were also removed per the manufacture’s recommendation. The cleaned reads (> 20bp) were then mapped to the mouse reference genome (GRCm38/mm10/ensemble release-84.38/ <u>GCA_000001635.6</u>) using STAR (30) (version 2.6.1a), allowing up to 2 mismatches depending on the alignment length (e.g. 20-29bp, 0 mismatches; 30-50bp, 1 mismatch; 50–60+bp, 2 mismatches). Reads mapping to > 20 locations were discarded. Gene level counts were quantified using HTSeq (htseq-counts) (37) (version 0.9.1) (mode: intersection-nonempty).

Genes with unique Entrez IDs and a minimum of ∼2 counts-per-million (CPM) in 4 or more samples were selected for statistical testing. This was followed by scaling normalization using the trimmed mean of M-values (TMM) method (38) to correct for compositional differences between sample libraries. Differential expression between naive and infected ears was evaluated using limma voomWithQualityWeights (39) with empirical bayes smoothing. Genes with Benjamini & Hochberg (40) adjusted p-values ≤ 0.05 and absolute fold-changes ≥ 1.5 were considered significant.

Gene Set Enrichment Analysis (GSEA) was carried out using Kyoto Encyclopedia of Genes and Genomes (KEGG) pathway databases and for each KEGG pathway, a p-value was calculated using hypergeometric test. Cut-off of both p < 0.05 and adjusted p-value/FDR value < 0.05 was applied to identify enriched KEGG pathways. DEGs that are more than 1.5-fold relative to controls were used as input, with upregulated and downregulated genes considered separately. Subsequently, the heat maps were generated using these genes with complex Heatmap. All analyses and visualizations were carried out using the statistical computing environment R version 3.6.3, RStudio version 1.2.5042, and Bioconductor version 3.11. The raw data from our bulk RNA-Seq analysis were deposited in Gene Expression Omnibus (GEO accession number— GSE273822).

### Ingenuity Pathway Analysis

To categorize the extensive list of differentially expressed genes identified by RNASeq, we performed Ingenuity Pathway Analysis (IPA). IPA allows for the upload and analyzation of high throughput data by placing the data into biological pathways, while also building networks to represent biological systems. To perform the IPA, we inputted our list of DEGs from the RNASeq data into the IPA software (Qiagen). We used a p-value cut-off of <0.05 so that anything below that would be considered for analysis. For the fold change (FC), we used a range of FC -2 to 2 so any values outside of that range would be analyzed by IPA.

### Statistical Analysis

Statistical analysis was performed using GraphPad Prism 9. Besides the scRNASeq of leishmanial lesions and total RNASeq of BMDMs where statistics are described above, a t-test was performed with *p* ≤ 0.05 being considered statistically significant. A Grubbs’ test was used to identify and mathematically remove outlier data points.

## Results

HIF-α signaling is a hallmark of lesions following *Leishmania* infection, but the specific cell types in lesions undergoing HIF-α activation are not known (11,12,27). To identify the cell types in leishmanial lesions that express the transcription factors HIF-1α and HIF-2α as well as their transcriptional target genes, we performed scRNASeq on lesions 4 weeks after dermal *L. major* inoculation (28). Specifically, single cells from the ears of infected and naive mice were bar-coded and sequenced using the droplet-based 10X Genomics Chromium platform. Unbiased hierarchical clustering using Seurat was performed to identify clusters indicative of individual cell types (Figure 1A). Of the 35 distinct cell types, HIF-1α is mainly expressed in keratinocytes, fibroblasts, chondrocytes, and endothelial cells in naïve uninfected skin. In contrast, HIF-1α is predominantly expressed by infiltrating cells including T cells, neutrophils, and monocyte-derived macrophages during *L. major* infection (Figure 1B). Of the 35 distinct cell types, HIF-2α is mainly expressed in fibroblasts, chondrocytes, and endothelial cells in naïve uninfected skin (Figure 1B). After infection, HIF-2α retains expression in fibroblasts, chondrocytes, and endothelial cells and is additionally expressed in infiltrating T cells and monocyte-derived macrophages during *L. major* infection (Figure 1B).

**Figure 1.**
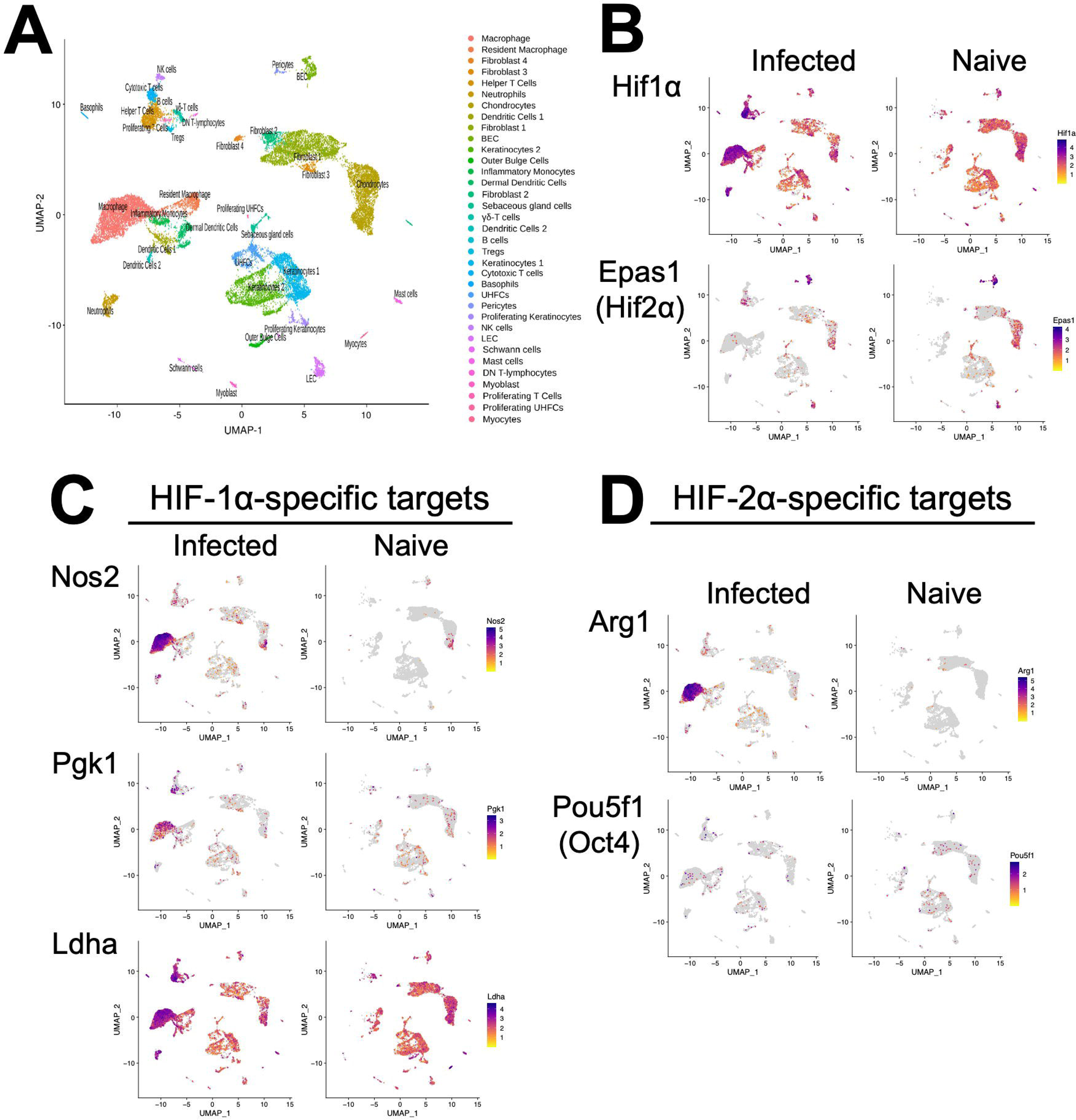
Single-cell RNASequencing shows HIF-α transcriptional targets are elevated in murine lesions during *L. major* infection. C57BL/6 mice were infected or not with 2×10^6^ *L. major* parasites intradermally in the ear. At 4 weeks infected ears and naïve uninfected control ears were digested and subjected to scRNASeq as a part of a previous study (28). (**A**) Uniform Manifold Approximation and Projection (UMAP) plot revealed 35 distinct cell clusters. Seurat’s FindClusters function identified each cell cluster and cell type designation to the right. To initially define cell clusters both naive and infected groups were combined, but here naive and infected groups are shown to specify transcript expression under each condition. (**B**) Feature plots of expression distribution for HIF-1α and HIF-2α (gene Epas1). (**C**) Feature plots of expression distribution for HIF-1α-specific target genes (Nos2, Pgk1, and Ldha). (**D**) Feature plots of expression distribution for HIF-2α-specific target genes (Arg1 and Oct4 (gene Pou5f1)). Expression levels for each gene are color-coded and overlaid onto UMAP plot. Cells with the highest expression level are colored dark purple.

Because HIF-α expression does not always correlate to HIF-α activity, we examined HIF-1α and HIF-2α transcriptional target genes as a surrogate for HIF-α activation. Besides Ldha, overall HIF-1α and HIF-2α target genes are expressed at low levels in naïve skin (Figure 1C-D). In contrast, HIF-1α-specific target genes including Nos2, Pgk1 and Ldha are dramatically increased upon infection, and these are predominantly expressed in monocyte-derived macrophages (Figure 1C). Similarly, the HIF-2α-specific target gene Arg1 is also highly expressed in monocyte-derived macrophages and Arg1 is significantly upregulated during infection (Figure 1D). However, another HIF-2α-specific target gene Pou5f1 (protein name Oct4) was only minorly expressed in monocyte-derived macrophages (Figure 1D). Altogether these transcriptomic data show that monocyte-derived macrophages exhibit HIF-1α and HIF-2α activation following *L. major* infection.

Given lesional monocyte-derived macrophages exhibited HIF-1α and HIF-2α activation, we evaluated the host macrophage responses during *L. major* infection using macrophages derived from monocytes from the bone marrow. To investigate the role of HIF-α signaling, we used macrophages from mice missing both HIF-1α and HIF-2α signaling where ARNT is deleted in myeloid cells and compared those responses to macrophages with intact HIF-1α and HIF-2α signaling (11,27,41). To first explore the host macrophage response in cells with intact HIF-1α and HIF-2α signaling, differential expression analysis was conducted on infected LysM^Cre^ARNT^f/+^ control macrophages compared to uninfected LysM^Cre^ARNT^f/+^ control macrophages referred to as ARNT^f/+^ going forward. Several differentially expressed genes (DEGs) were upregulated with *L. major* infection including *Gm15564*, *Gca, Stk35*, and *Socs1* while only *Tcf4* was found to be downregulated with *L. major* infection (Figure 2A and Table 1). KEGG analysis revealed several enriched pathways with *L. major* infection including the ‘PPAR signaling pathway’, ‘Rap1 signaling pathway’, and ‘Chemokine signaling pathway’ (Figure 2C). Additionally, the ‘IL-17 cell differentiation pathway’ was downregulated in infected macrophages compared to uninfected ARNT^f/+^ macrophages (Figure 2C). Of note, the ‘HIF-1α signaling pathway’ was upregulated with infection (Figure 2C). These data demonstrate that infection with *L. major* is sufficient to drive transcriptional changes in macrophages and activate HIF-α signaling.

**Figure 2:**
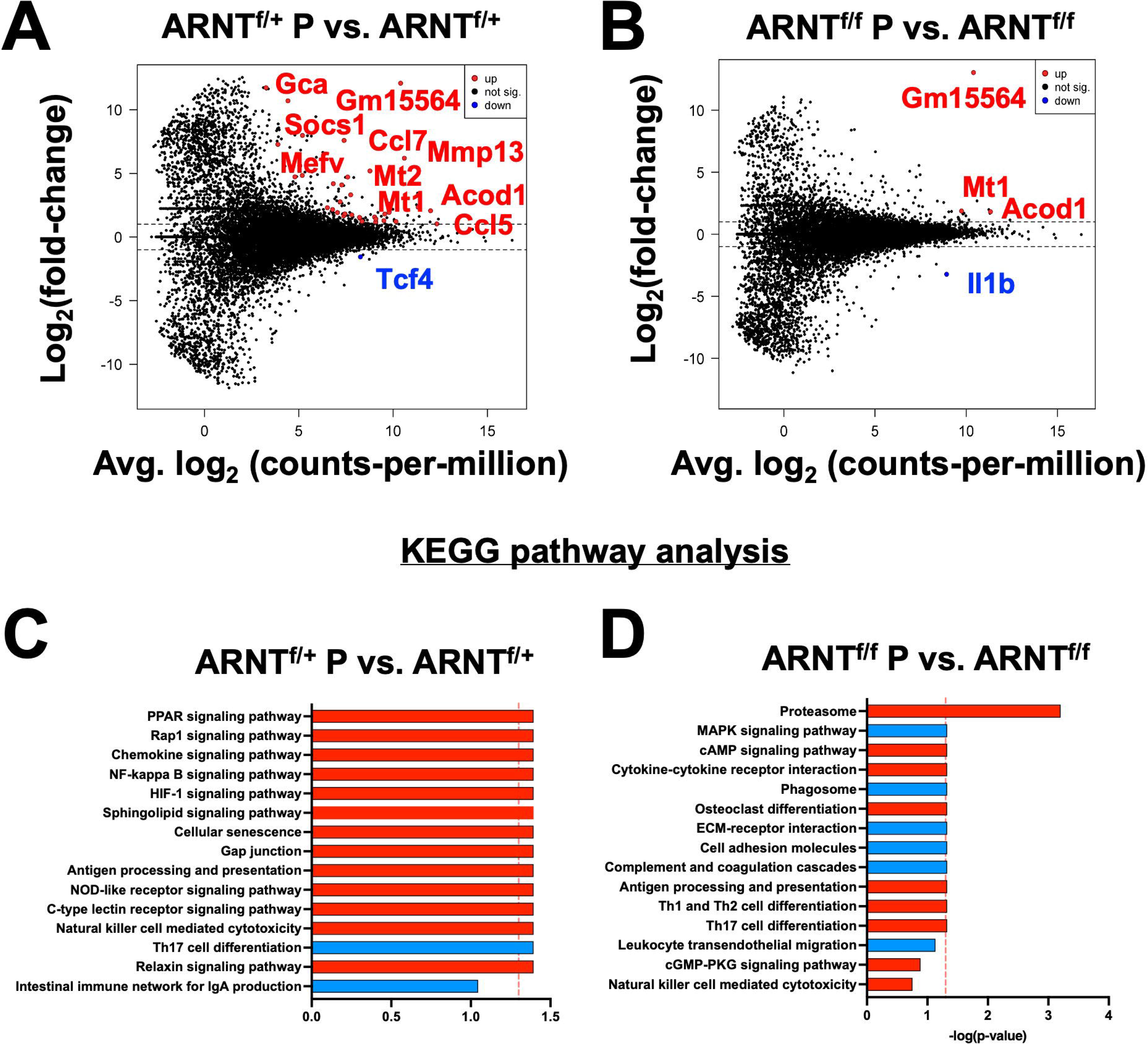
Infection with *L. major* induces transcriptional changes that are absent in infected macrophages deficient for HIF-α signaling. Macrophages infected with *L. major* were lysed and prepped for RNASequencing. (**A**) A mean-difference plot (MD plot) depicts transcripts upregulated (red) and downregulated (blue) with infection in macrophages with intact HIF-α signaling (ARNT^f/+^ P vs. ARNT^f/+^) where P indicates parasites. (**B**) An MD plot shows upregulated and downregulated transcripts in infected macrophages deficient for HIF-α signaling (ARNT^f/f^ P vs. ARNT^f/f^). (**C**) KEGG analysis was performed to identify the top enriched pathways during infection with *L. major* in macrophages with or without HIF-α signaling. Upregulated pathways are shown in red and downregulated pathways are shown in blue.

**Table 1.** Significantly Up- or Down-regulated DEGs between ARNT^f/+^ Infected Macrophages Compared to ARNT^f/+^ Uninfected Macrophages.

| Up-regulated | GENE NAME | logFC | PValue (adj.) |
| --- | --- | --- | --- |
| SYMBOL |  |  |  |
| Gm15564 | Predicted gene 15564 | 12.1 | 5.93E-05 |
| Gca | grancalcin | 12.0 | 0.00541 |
| Stk35 | serine/threonine kinase 35 | 11.7 | 0.007409 |
| Socs1 | suppressor of cytokine signaling 1 | 10.7 | 0.003167 |
| Xkr8 | X-linked Kx blood group related 8 | 7.99 | 0.024124 |
| Ccl7 | chemokine (C-C motif) ligand 7 | 7.60 | 1.39E-10 |
| Eaf1 | ELL associated factor 1 | 7.28 | 0.027228 |
| Mmp13 | matrix metalloproteinase 13 | 6.19 | 0.009873 |
| Mefv | Mediterranean fever | 5.61 | 0.054491 |
| Mt2 | grancalcin | 5.20 | 0.001488 |
| Gsr | glutathione reductase | 4.84 | 2.41E-08 |
| Fkbp2 | FK506 binding protein 2 | 4.72 | 0.050311 |
| Gnb4 | guanine nucleotide binding protein (G protein), beta 4 | 4.72 | 0.007542 |
| Slfn1 | schlafen 1 | 4.19 | 0.000198 |
| Fpr2 | formyl peptide receptor 2 | 4.09 | 0.017288 |
| Isg20 | interferon-stimulated protein | 3.32 | 0.021429 |
| Mmp12 | matrix metalloproteinase 12 | 2.76 | 1.39E-10 |
| Mt1 | metallothionein 1 | 2.60 | 0.018192 |
| Acod1 | aconitate decarboxylase 1 | 2.35 | 0.007409 |
| Tent5c | terminal nucleotidyltransferase 5C | 2.31 | 0.00651 |
| Mcoln2 | mucolipin 2 | 2.16 | 0.005174 |
| Il1rn | interleukin 1 receptor antagonist | 2.06 | 0.003563 |
| Clec4e | C-type lectin domain family 4, member e | 1.93 | 0.027228 |
| Kmt5a | lysine methyltransferase 5A | 1.93 | 0.00541 |
| Selenos | selenoprotein S | 1.82 | 1.59E-08 |
| Down-regulated |  |  |  |
| Symbol | GENE NAME | logFC | PValue (adj.) |
| Tcf4 | transcription factor 4 | -1.56 | 0.005174 |

We next investigated transcriptional changes during *L. major* infection in macrophages devoid of HIF-α signaling by comparing the transcriptome of infected LysM^Cre^ARNT^f/f^ macrophages to uninfected LysM^Cre^ARNT^f/f^ macrophages referred to as ARNT^f/f^ for the duration of the study. Four genes were differentially expressed, including *Mt1*, *Acod1*, *Il1β* and a predicted gene, *gm15564* (Figure 2B and Table 2). *Mt1, Acod1,* and *gm15564* were also upregulated with infection in HIF-α competent macrophages, indicating these transcriptional changes are independent of HIF-α signaling (Figure 2A and Table 1). Most of the transcriptomic changes seen during *L. major* infection were ablated in the absence of HIF-α signaling. For instance, 22 DEGs were upregulated in HIF-α competent macrophages with infection, that were not detected during infection in macrophages deficient for HIF-α signaling (Figure 2A and Table 1). To further characterize the cellular processes most affected by infection in macrophages either with or without HIF-α signaling, we conducted KEGG pathway analyses. The analysis revealed that during infection with *L. major,* the proteasome pathway is upregulated in the absence of HIF-α signaling (Figure 2D). When we investigated enriched pathways in infected macrophages with intact HIF-α, the proteosome pathway was not upregulated suggesting this pathway is normally suppressed by HIF-α during *L. major* infection. These data together indicate infection alone induces transcriptional changes that are HIF-α-dependent, suggesting infection with *L. major* parasites is sufficient to activate HIF-α signaling in vitro contrary to other reports (23,25,27).

**Table 2.**
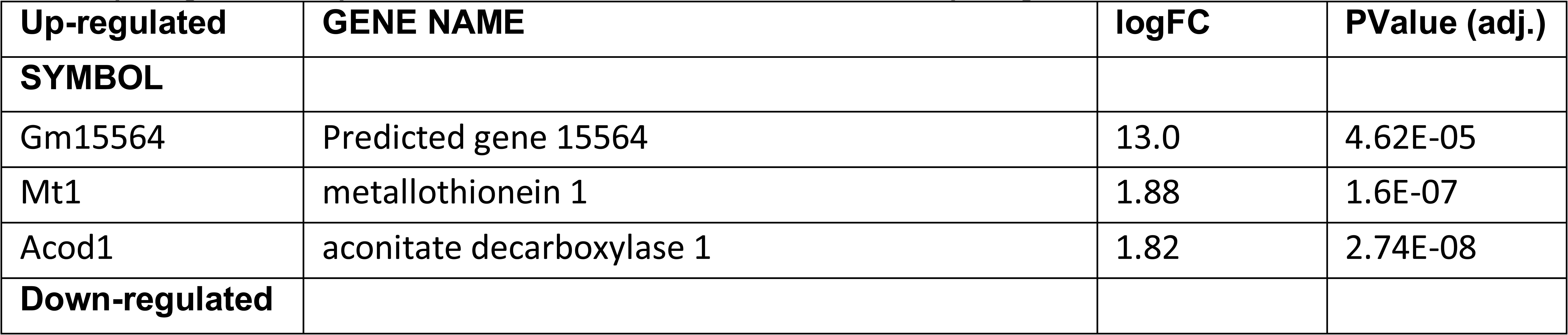
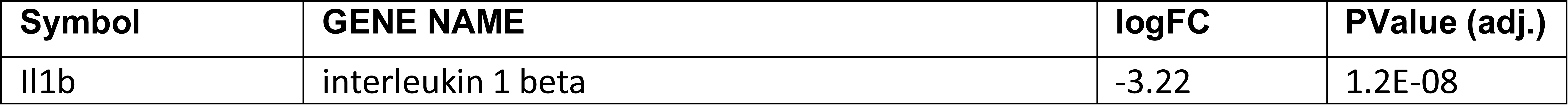
Significantly Up- or Down-regulated DEGs between ARNT^f/f^ Infected Macrophages Compared to ARNT^f/f^ Uninfected Macrophages.

During infection with *L. major* parasites, a strong Th1 immune response is formed resulting in the release of a multitude of pro-inflammatory mediators. To identify inflammation-related transcriptomic changes, both HIF-α signaling competent and deficient macrophages were infected with *L. major* and treated with LPS and IFNγ to mimic the in vivo pro-inflammatory environment. We identified 1,076 genes that were differentially expressed when comparing infected ARNT^f/+^ macrophages treated with LPS and IFNγ to infected ARNT^f/+^ macrophages not treated with LPS and IFNγ (Figure 3A and Table 3). The top upregulated DEGs were *Gpr18* and *Mmp25* (Table 3). Additionally, there were many immune-related transcripts that were upregulated to a lesser extent including *Il12b*, *Cd40*, and *Nos2*, all of which participate in the immune response to *Leishmania* parasites (Figure 3A and Table 3). The top downregulated DEGs were *Arrdc3*, *Rasgrp3*, and *Cdca7l* comparing infected ARNT^f/+^ macrophages treated or not with LPS and IFNγ (Table 3). Furthermore, we investigated transcriptional changes in infected ARNT^f/f^ macrophages treated with LPS and IFNγ compared to infected ARNT^f/f^ not treated with LPS and IFNγ. We identified 1,191 DEGs (Figure 3B and Table 4). The top 25 most upregulated genes in response to pro-inflammatory stimuli were the same for infected macrophages with or without competent HIF-α signaling, indicating these DEGs are independent of HIF-α signaling (Figure 3A-B and Table 4). In contrast, there were differences in the top 25 downregulated DEGs in response to LPS and IFNγ stimulation in infected macrophages that are competent or impaired for HIF-α signaling (Figure 3B and Table 4). The top downregulated DEGs in infected HIF-α deficient macrophages treated with LPS and IFNγ compared to infected HIF-α deficient macrophages not treated with LPS and IFNγ were *Mdp1*, *Arap3*, and *Prmt3* (Table 4).

**Figure 3:**
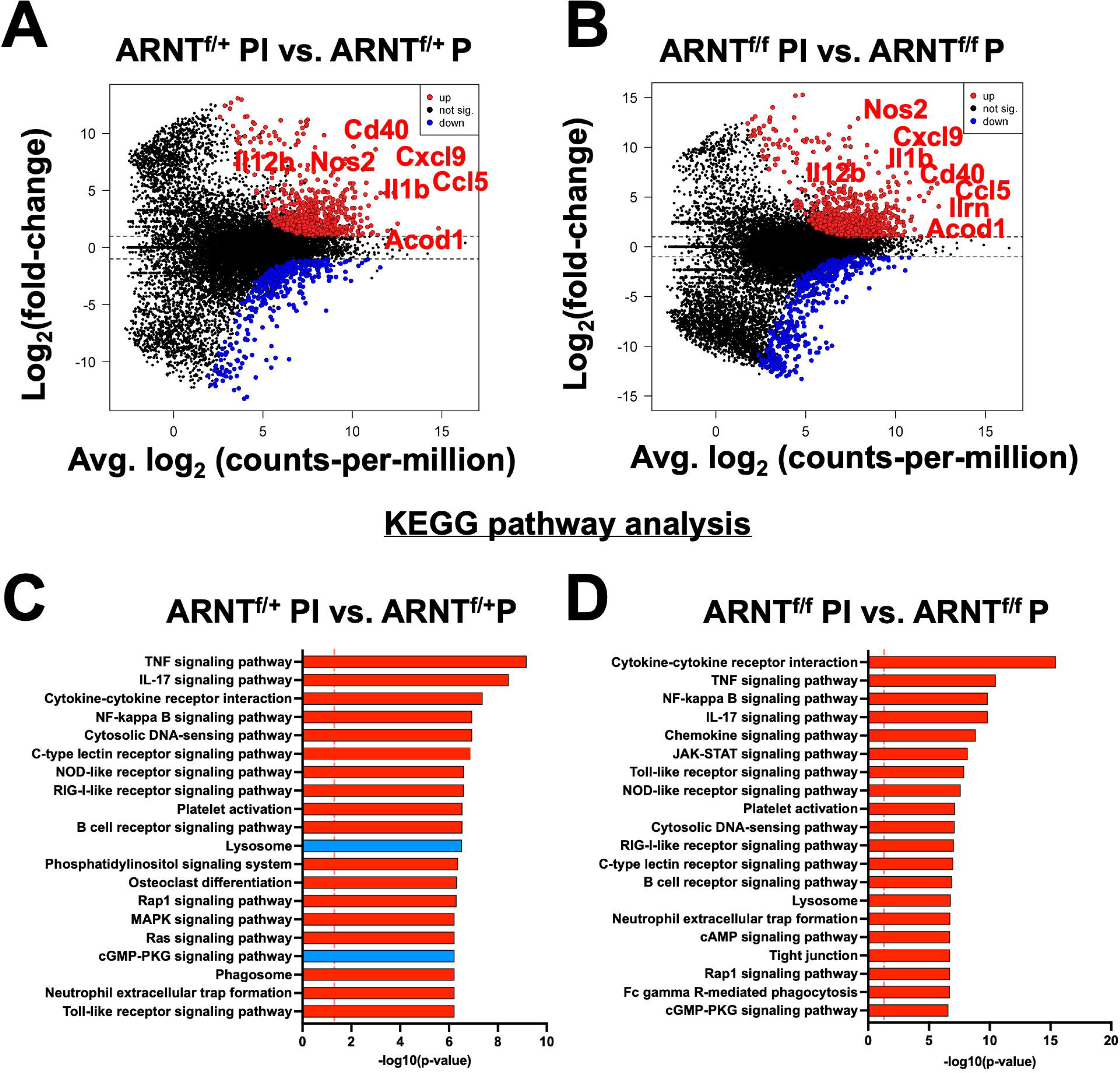
Inflammation related transcriptomic changes are both HIF-α dependent and independent. Macrophages were infected or infected and treated with LPS/IFNy and prepped for subsequent analysis utilizing RNASeq. (**A-B**) An MD plot illustrates transcriptomic changes during infection and stimulation with LPS/IFNy in HIF-α competent (ARNT^f/+^ PI vs. ARNT^f/+^P) and HIF-α deficient macrophages (ARNT^f/f^ PI vs. ARNT^f/f^ P). Here the PI indicates parasites and inflammatory stimuli, LPS/IFNy, and P describes parasite infection alone. Red dots identify upregulated transcripts and blue dots identify downregulated transcripts. (**C-D**) Enriched pathways were identified using KEGG analysis for both comparisons. Red pathways indicate upregulation while blue pathways are downregulated.

**Table 3.**
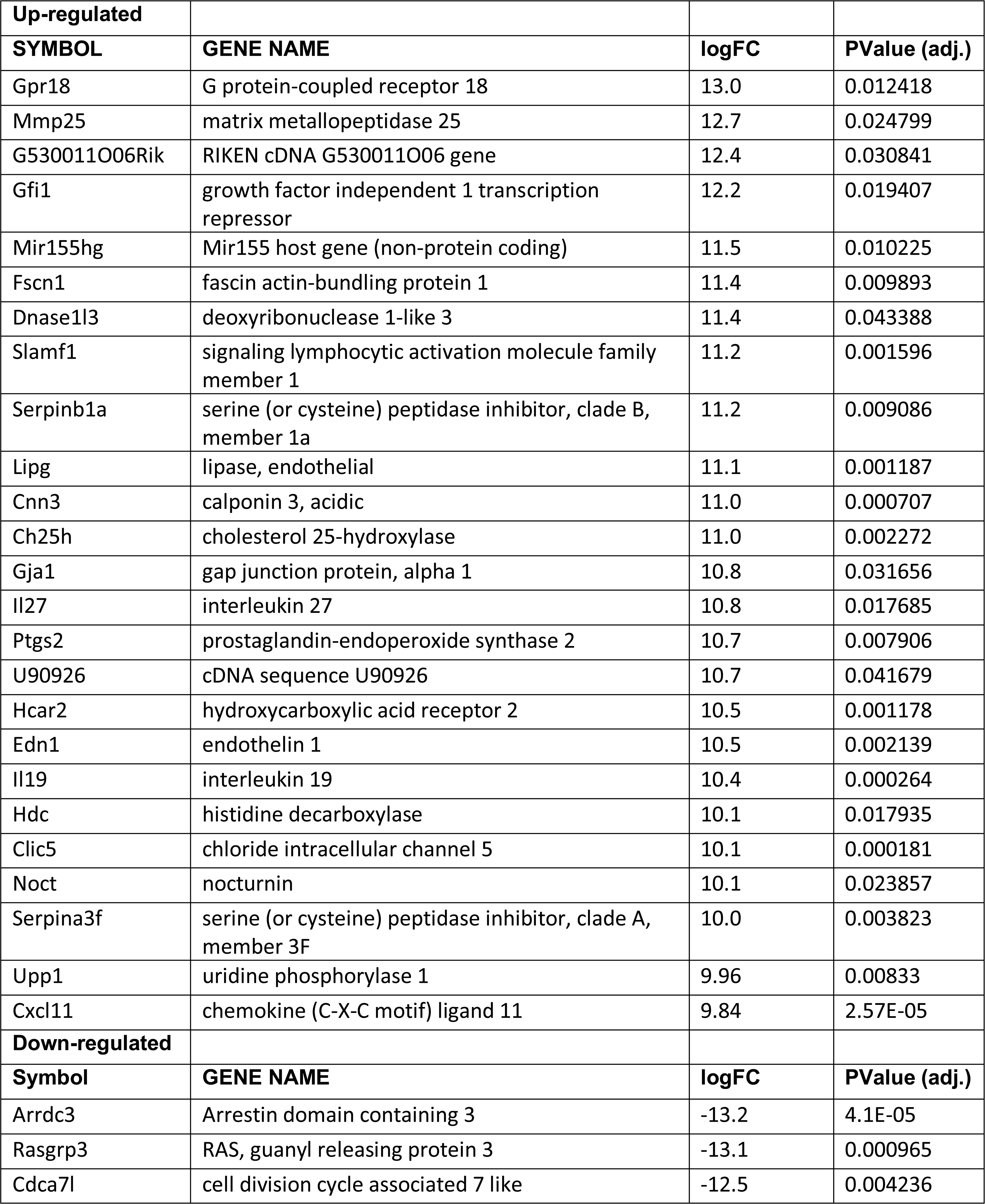
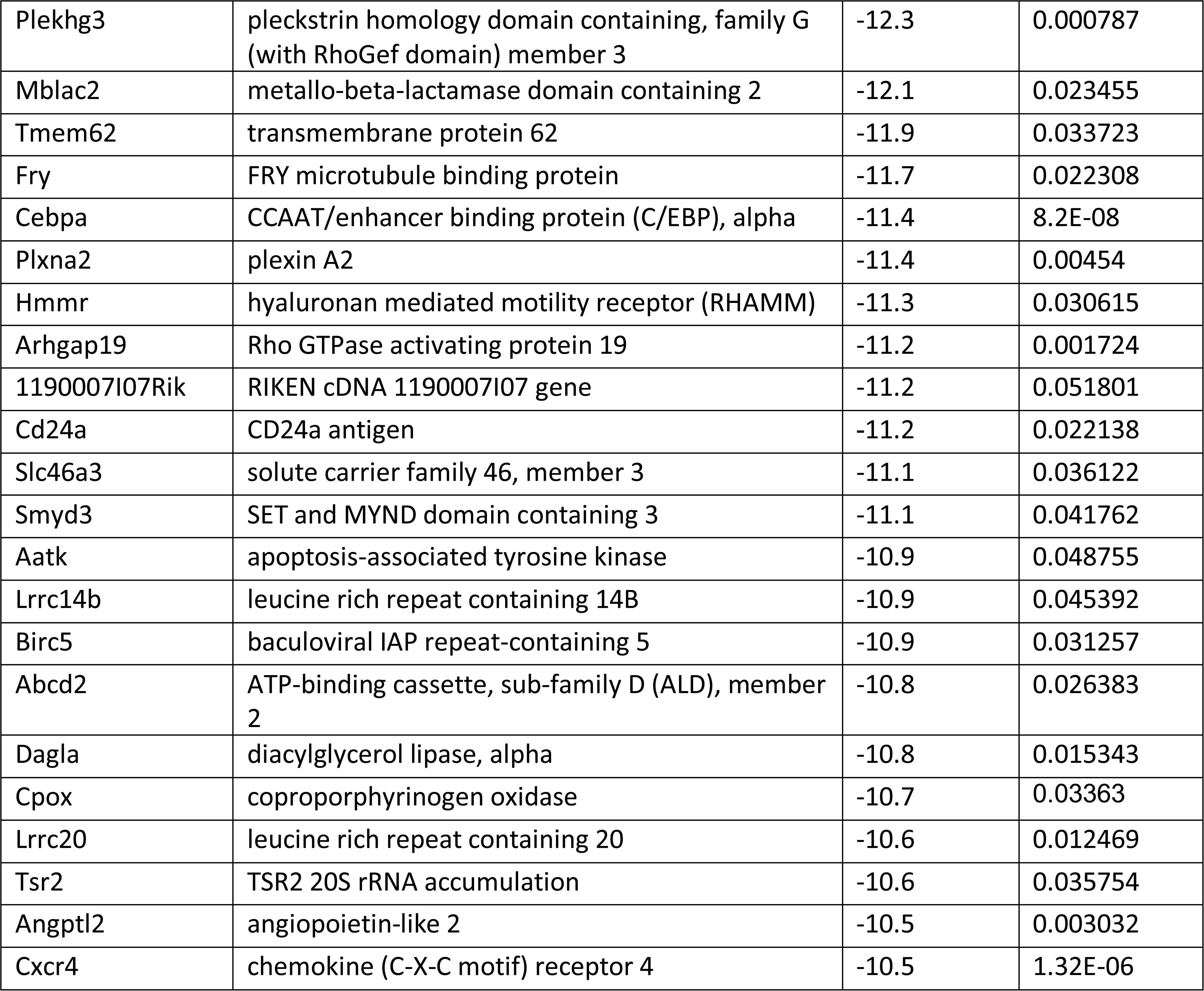
Significantly Up- or Down-regulated DEGs between Infected ARNT^f/+^ Macrophages Treated with LPS/ IFNg Compared to Infected ARNT^f/+^ Macrophages.

| Up-regulated |  |  |  |
| --- | --- | --- | --- |
| SYMBOL | GENE NAME | logFC | PValue (adj.) |
| Gpr18 | G protein-coupled receptor 18 | 13.0 | 0.012418 |
| Mmp25 | matrix metalloproteinase 25 | 12.7 | 0.024799 |
| G530011O06Rik | RIKEN cDNA G530011O06 gene | 12.4 | 0.030841 |
| Gfi1 | growth factor independent 1 transcription repressor | 12.2 | 0.019407 |
| Mir155hg | Mir155 host gene (non-protein coding) | 11.5 | 0.010225 |
| Fscn1 | fascin actin-bundling protein 1 | 11.4 | 0.009893 |
| Dnase1l3 | deoxyribonuclease 1-like 3 | 11.4 | 0.043388 |
| Slamf1 | signaling lymphocytic activation molecule family member 1 | 11.2 | 0.001596 |
| Serpinb1a | serine (or cysteine) peptidase inhibitor, clade B, member 1a | 11.2 | 0.009086 |
| Lipg | lipase, endothelial | 11.1 | 0.001187 |
| Cnn3 | calponin 3, acidic | 11.0 | 0.000707 |
| Ch25h | cholesterol 25-hydroxylase | 11.0 | 0.002272 |
| Gja1 | gap junction protein, alpha 1 | 10.8 | 0.031656 |
| Il27 | interleukin 27 | 10.8 | 0.017685 |
| Ptgs2 | prostaglandin-endoperoxide synthase 2 | 10.7 | 0.007906 |
| U90926 | cDNA sequence U90926 | 10.7 | 0.041679 |
| Hcar2 | hydroxycarboxylic acid receptor 2 | 10.5 | 0.001178 |
| Edn1 | endothelin 1 | 10.5 | 0.002139 |
| Il19 | interleukin 19 | 10.4 | 0.000264 |
| Hdc | histidine decarboxylase | 10.1 | 0.017935 |
| Clic5 | chloride intracellular channel 5 | 10.1 | 0.000181 |
| Noct | nocturnin | 10.1 | 0.023857 |
| Serpina3f | serine (or cysteine) peptidase inhibitor, clade A, member 3F | 10.0 | 0.003823 |
| Upp1 | uridine phosphorylase 1 | 9.96 | 0.00833 |
| Cxcl11 | chemokine (C-X-C motif) ligand 11 | 9.84 | 2.57E-05 |
| Down-regulated |  |  |  |
| Symbol | GENE NAME | logFC | PValue (adj.) |
| Arrdc3 | Arrestin domain containing 3 | -13.2 | 4.1E-05 |
| Rasgrp3 | RAS, guanyl releasing protein 3 | -13.1 | 0.000965 |
| Cdca7l | cell division cycle associated 7 like | -12.5 | 0.004236 |
| Plekhg3 | pleckstrin homology domain containing, family G (with RhoGef domain) member 3 | -12.3 | 0.000787 |
| Mblac2 | metallo-beta-lactamase domain containing 2 | -12.1 | 0.023455 |
| Tmem62 | transmembrane protein 62 | -11.9 | 0.033723 |
| Fry | FRY microtubule binding protein | -11.7 | 0.022308 |
| Cebpa | CCAAT/enhancer binding protein (C/EBP), alpha | -11.4 | 8.2E-08 |
| Plxna2 | plexin A2 | -11.4 | 0.00454 |
| Hmmr | hyaluronan mediated motility receptor (RHAMM) | -11.3 | 0.030615 |
| Arhgap19 | Rho GTPase activating protein 19 | -11.2 | 0.001724 |
| 1190007I07Rik | RIKEN cDNA 1190007I07 gene | -11.2 | 0.051801 |
| Cd24a | CD24a antigen | -11.2 | 0.022138 |
| Slc46a3 | solute carrier family 46, member 3 | -11.1 | 0.036122 |
| Smyd3 | SET and MYND domain containing 3 | -11.1 | 0.041762 |
| Aatk | apoptosis-associated tyrosine kinase | -10.9 | 0.048755 |
| Lrrc14b | leucine rich repeat containing 14B | -10.9 | 0.045392 |
| Birc5 | baculoviral IAP repeat-containing 5 | -10.9 | 0.031257 |
| Abcd2 | ATP-binding cassette, sub-family D (ALD), member 2 | -10.8 | 0.026383 |
| Dagla | diacylglycerol lipase, alpha | -10.8 | 0.015343 |
| Cpox | coproporphyrinogen oxidase | -10.7 | 0.03363 |
| Lrrc20 | leucine rich repeat containing 20 | -10.6 | 0.012469 |
| Tsr2 | TSR2 20S rRNA accumulation | -10.6 | 0.035754 |
| Angptl2 | angiopoietin-like 2 | -10.5 | 0.003032 |
| Cxcr4 | chemokine (C-X-C motif) receptor 4 | -10.5 | 1.32E-06 |

**Table 4.**
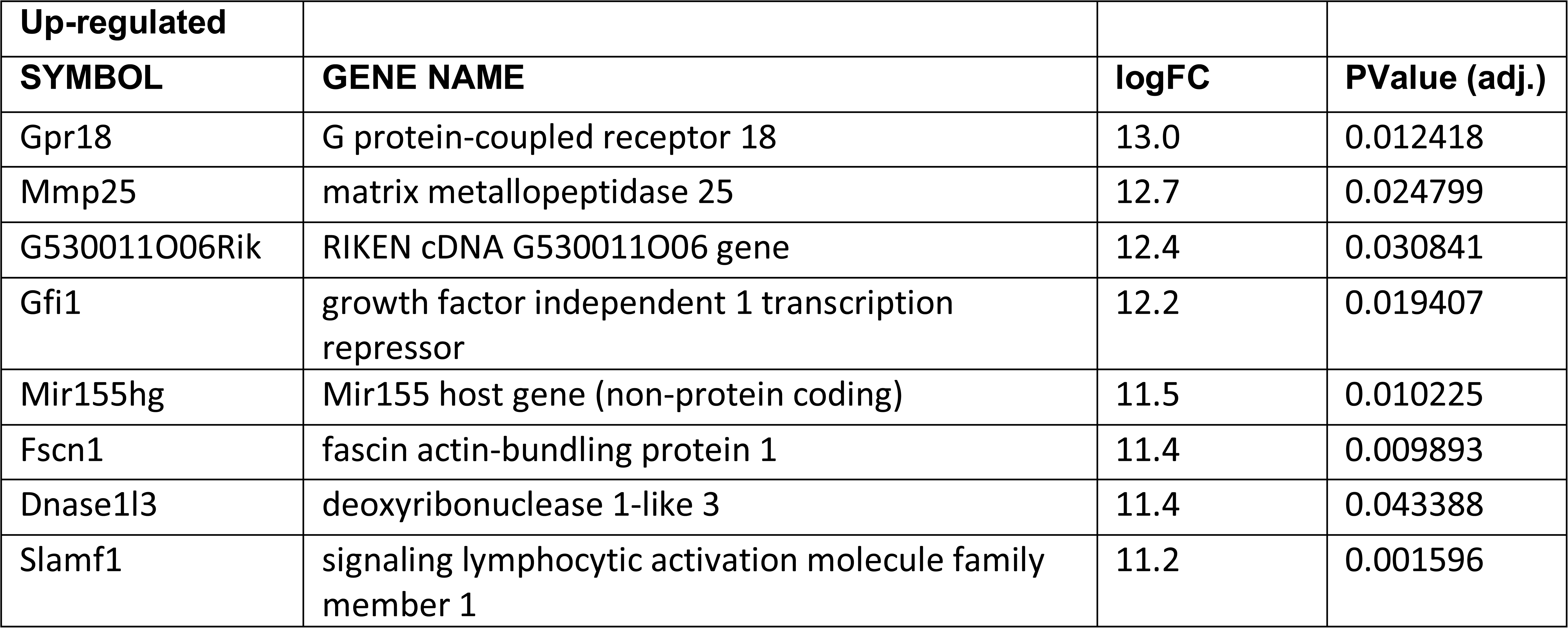
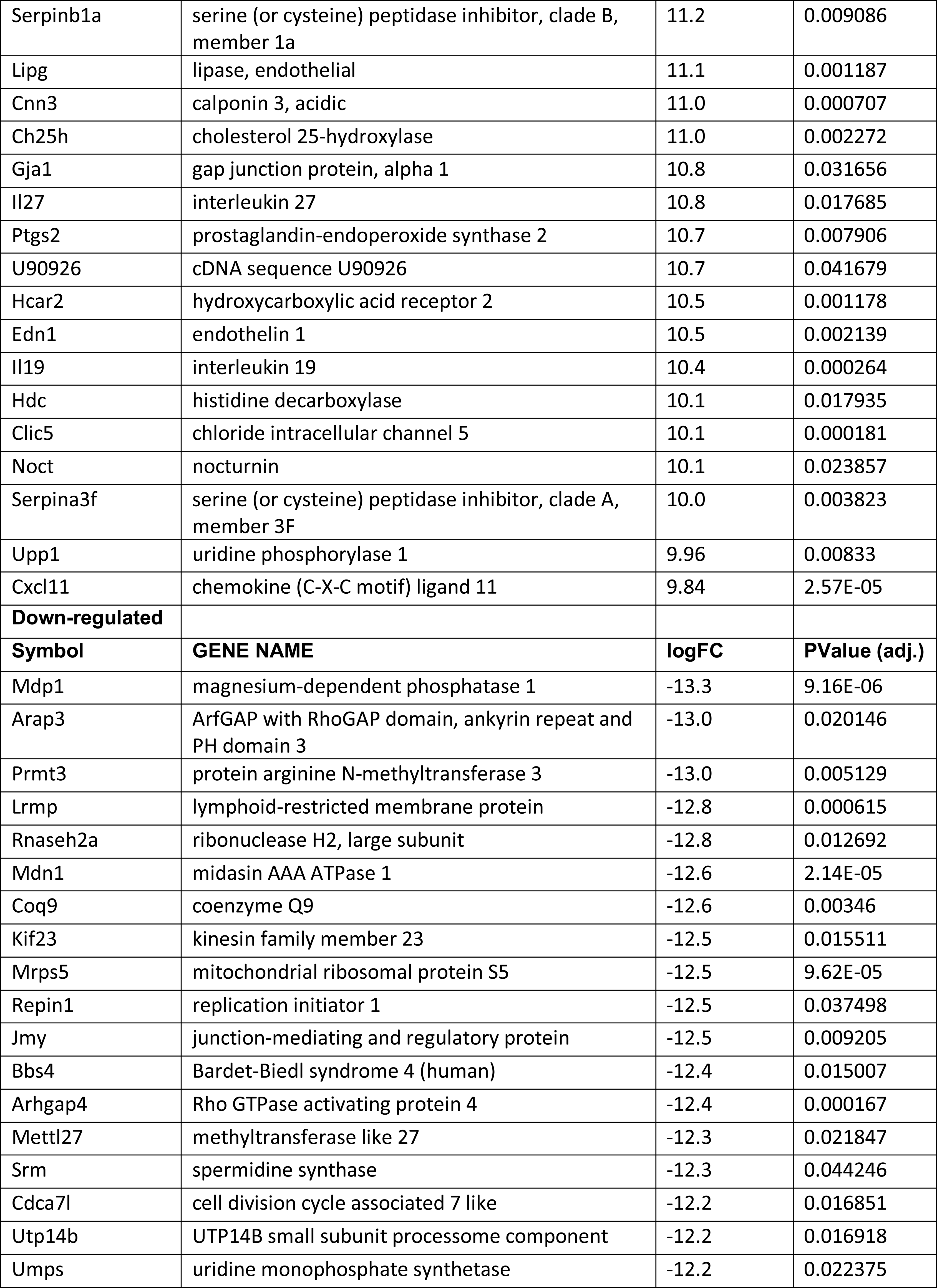
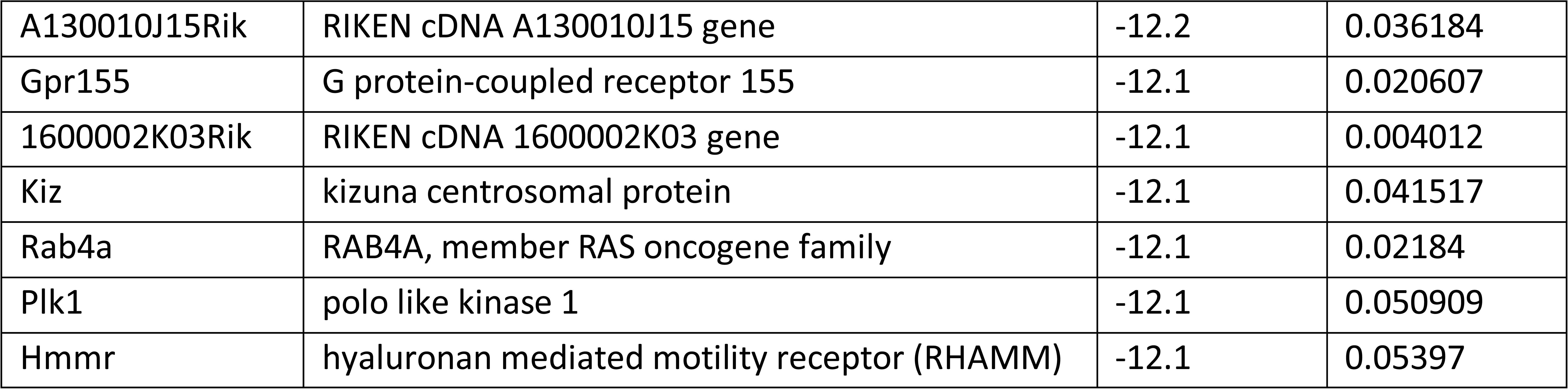
Significantly Up- or Down-regulated DEGs between Infected ARNT^f/f^ Macrophages Treated with LPS/ IFNg Compared to Infected ARNT^f/f^ Macrophages.

| Up-regulated |  |  |  |
| --- | --- | --- | --- |
| SYMBOL | GENE NAME | logFC | PValue (adj.) |
| Gpr18 | G protein-coupled receptor 18 | 13.0 | 0.012418 |
| Mmp25 | matrix metalloproteinase 25 | 12.7 | 0.024799 |
| G530011O06Rik | RIKEN cDNA G530011O06 gene | 12.4 | 0.030841 |
| Gfi1 | growth factor independent 1 transcription repressor | 12.2 | 0.019407 |
| Mir155hg | Mir155 host gene (non-protein coding) | 11.5 | 0.010225 |
| Fscn1 | fascin actin-bundling protein 1 | 11.4 | 0.009893 |
| Dnase1l3 | deoxyribonuclease 1-like 3 | 11.4 | 0.043388 |
| Slamf1 | signaling lymphocytic activation molecule family member 1 | 11.2 | 0.001596 |
| Serpinb1a | serine (or cysteine) peptidase inhibitor, clade B, member 1a | 11.2 | 0.009086 |
| Lipg | lipase, endothelial | 11.1 | 0.001187 |
| Cnn3 | calponin 3, acidic | 11.0 | 0.000707 |
| Ch25h | cholesterol 25-hydroxylase | 11.0 | 0.002272 |
| Gja1 | gap junction protein, alpha 1 | 10.8 | 0.031656 |
| Il27 | interleukin 27 | 10.8 | 0.017685 |
| Ptgs2 | prostaglandin-endoperoxide synthase 2 | 10.7 | 0.007906 |
| U90926 | cDNA sequence U90926 | 10.7 | 0.041679 |
| Hcar2 | hydroxycarboxylic acid receptor 2 | 10.5 | 0.001178 |
| Edn1 | endothelin 1 | 10.5 | 0.002139 |
| Il19 | interleukin 19 | 10.4 | 0.000264 |
| Hdc | histidine decarboxylase | 10.1 | 0.017935 |
| Clic5 | chloride intracellular channel 5 | 10.1 | 0.000181 |
| Noct | nocturnin | 10.1 | 0.023857 |
| Serpina3f | serine (or cysteine) peptidase inhibitor, clade A, member 3F | 10.0 | 0.003823 |
| Upp1 | uridine phosphorylase 1 | 9.96 | 0.00833 |
| Cxcl11 | chemokine (C-X-C motif) ligand 11 | 9.84 | 2.57E-05 |
| **Down-regulated** |  |  |  |
| **Symbol** | **GENE NAME** | **logFC** | **PValue (adj.)** |
| Mdp1 | magnesium-dependent phosphatase 1 | -13.3 | 9.16E-06 |
| Arap3 | ArfGAP with RhoGAP domain, ankyrin repeat and PH domain 3 | -13.0 | 0.020146 |
| Prmt3 | protein arginine N-methyltransferase 3 | -13.0 | 0.005129 |
| Lrmp | lymphoid-restricted membrane protein | -12.8 | 0.000615 |
| Rnaseh2a | ribonuclease H2, large subunit | -12.8 | 0.012692 |
| Mdn1 | midasin AAA ATPase 1 | -12.6 | 2.14E-05 |
| Coq9 | coenzyme Q9 | -12.6 | 0.00346 |
| Kif23 | kinesin family member 23 | -12.5 | 0.015511 |
| Mrps5 | mitochondrial ribosomal protein S5 | -12.5 | 9.62E-05 |
| Repin1 | replication initiator 1 | -12.5 | 0.037498 |
| Jmy | junction-mediating and regulatory protein | -12.5 | 0.009205 |
| Bbs4 | Bardet-Biedl syndrome 4 (human) | -12.4 | 0.015007 |
| Arhgap4 | Rho GTPase activating protein 4 | -12.4 | 0.000167 |
| Mettl27 | methyltransferase like 27 | -12.3 | 0.021847 |
| Srm | spermidine synthase | -12.3 | 0.044246 |
| Cdca7l | cell division cycle associated 7 like | -12.2 | 0.016851 |
| Utp14b | UTP14B small subunit processome component | -12.2 | 0.016918 |
| Umps | uridine monophosphate synthetase | -12.2 | 0.022375 |
| A130010J15Rik | RIKEN cDNA A130010J15 gene | -12.2 | 0.036184 |
| Gpr155 | G protein-coupled receptor 155 | -12.1 | 0.020607 |
| 1600002K03Rik | RIKEN cDNA 1600002K03 gene | -12.1 | 0.004012 |
| Kiz | kizuna centrosomal protein | -12.1 | 0.041517 |
| Rab4a | RAB4A, member RAS oncogene family | -12.1 | 0.02184 |
| Plk1 | polo like kinase 1 | -12.1 | 0.050909 |
| Hmmr | hyaluronan mediated motility receptor (RHAMM) | -12.1 | 0.05397 |

A functional analysis was performed to identify pathways associated with pro-inflammatory stimulus administration in infected macrophages with or without HIF-α signaling compared to their infected macrophage counterparts with no stimulus. The KEGG analysis revealed pro-inflammatory stimuli upregulated pathways such as ‘TNF signaling receptor’, ‘IL-17 signaling pathway’, and several other inflammatory pathways (Figure 3C). In addition, these infected macrophages downregulated the ‘lysosome’ and ‘cGMP-PKG pathway’ in response to LPS and IFNγ administration (Figure 3C). Next, we compared infected ARNT^f/f^ macrophages treated or not with LPS and IFNγ by KEGG analysis. The results revealed similar upregulated pathways as the pro-inflammatory treated and infected ARNT^f/+^ macrophages including ‘cytokine-cytokine receptor interaction’ and ‘TNF signaling pathway’ (Figure 3D). Interestingly, there were no significantly downregulated pathways in infected pro-inflammatory stimulated macrophages deficient for HIF-α signaling compared to infected macrophages also deficient for HIF-α signaling. In contrast to the HIF-α competent macrophages, during HIF-α deficiency, the ‘lysosome’ and ‘cGMP-Pk3 signaling pathways’ were upregulated in infected macrophages treated with LPS and IFNγ (Figure 3D). These data suggest in a pro-inflammatory environment, HIF-α suppresses pathways related to *L. major* infection including those involved in production of the phagolysosome and second messenger signaling.

After investigating changes involved with *L. major* infection alone and with pro-inflammatory stimulus in macrophages with and without competent HIF-α signaling, we directly compared the gene expression profiles of macrophages without HIF-α signaling to macrophages with HIF-α signaling under each condition. First, we compared ARNT^f/f^ to ARNT^f/+^ under basal conditions. We identified two upregulated DEGs in the ARNT^f/f^ macrophages compared to ARNT^f/+^ macrophages including, *Isg20* and *Spp1* suggesting HIF-α inhibits these genes during homeostasis (Figure 4A and Table 5). Next, we analyzed differences between ARNT^f/f^ to ARNT^f/+^ during infection with *L. major* parasites. When comparing macrophages without or with HIF-α signaling during *L. major* infection, we found several downregulated DEGs in macrophages with impaired HIF-α signaling which suggests under normal conditions these DEGs are mediated by HIF-α (Figure 4B and Table 6). These DEGs include *Il1β*, *Ccl5*, *Mcoln2*, *Mevf*, and *Socs1* (Figure 4B and Table 6). By KEGG analysis, we found under basal conditions macrophages without HIF-α signaling upregulate the ‘ribosome’ and ‘DNA replication pathways’ compared to macrophages with HIF-α signaling (Figure 4C). This finding suggests that HIF-α restricts cell processes in the absence of infection in steady state.

**Figure 4:**
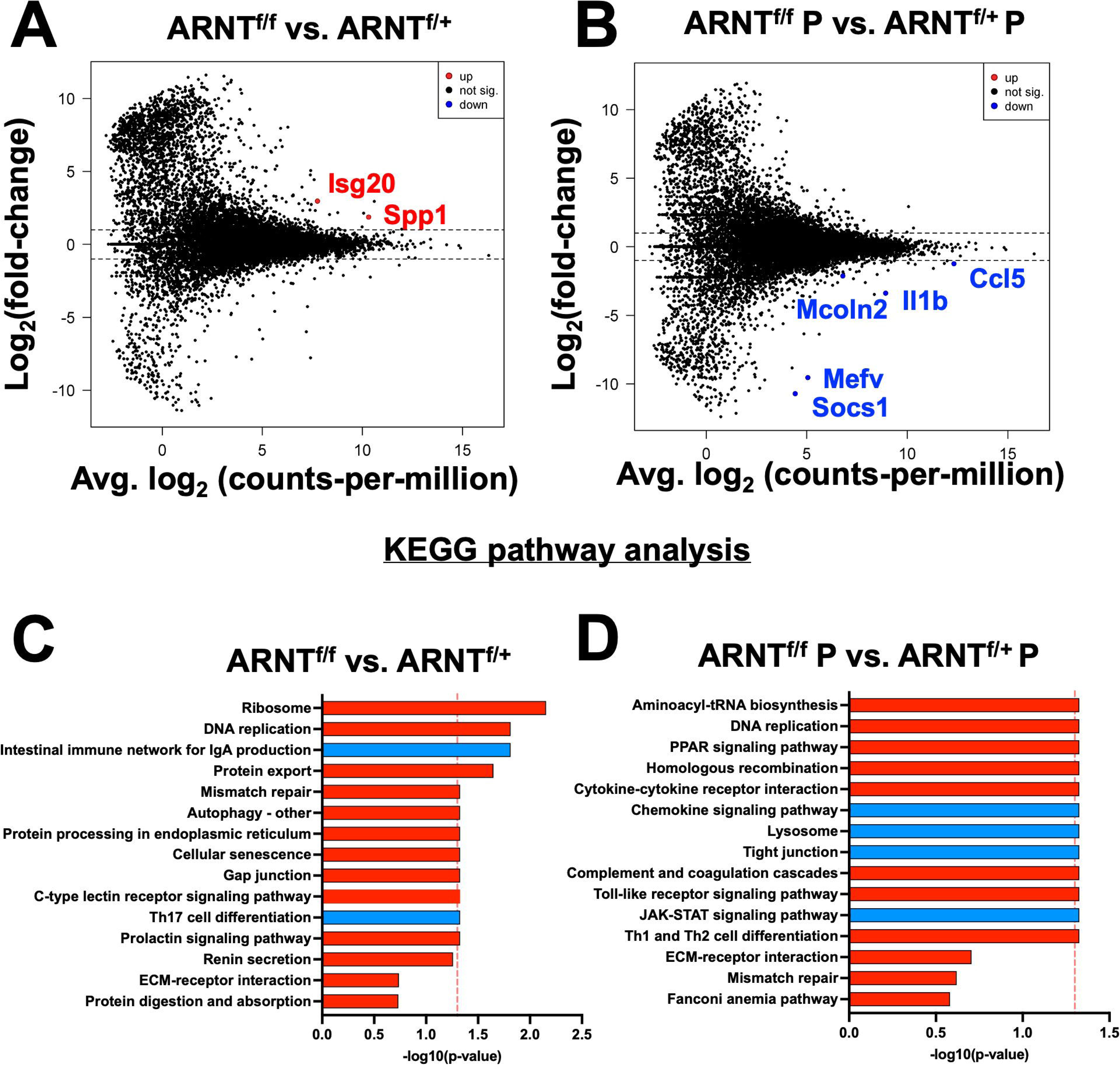
HIF-α mediates DEGs induced by *L. major* parasites. Macrophages both with and without HIF-a signaling were cultured in media alone or infected with *L. major* parasites for 8 hours before being prepped for RNASequencing. (**A**) An MD plot illustrates transcripts inhibited by HIF-α under basal conditions. (**B**) An MD plot shows transcripts mediated by HIF-α during *L. major* infection. (**C**) KEGG pathway analysis identified enriched pathways in macrophages without HIF-α signaling under basal conditions. (**D**) KEGG pathway analysis identified enriched pathways in macrophages without HIF-α signaling during infection. Red pathways are upregulated and blue pathways are downregulated.

**Table 5.**
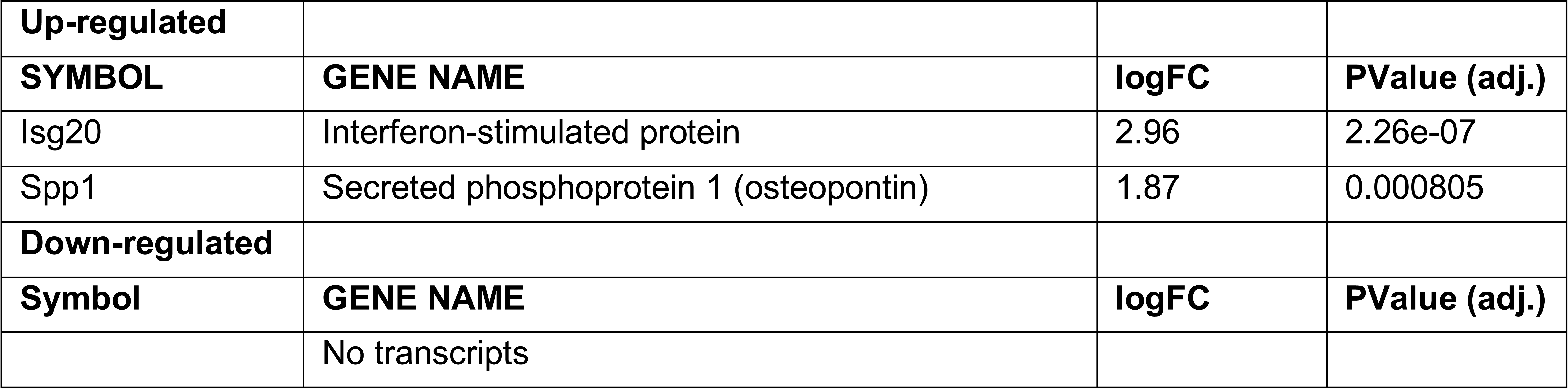
Significantly Up- or Down-regulated DEGs between ARNT^f/f^ compared to ARNT^f/+^ Uninfected Macrophages.

**Table 6.** Significantly Up- or Down-regulated DEGs between ARNT^f/f^ compared to ARNT^f/+^ Infected Macrophages.

|  |  |  |  |
| --- | --- | --- | --- |
| **Up-regulated** |  |  |  |
| **SYMBOL** | **GENE NAME** | **logFC** | **PValue (adj.)** |
|  | No transcripts |  |  |
| **Down-regulated** |  |  |  |
| **Symbol** | **GENE NAME** | **logFC** | **PValue (adj.)** |
| Socs1 | Suppressor of cytokine signaling 1 | -10.7 | 0.050445 |
| Mefv | Mediterranean fever | -9.56 | 0.00394 |
| Il1b | Interleukin 1 beta | -3.38 | 1.36e-08 |
| Mcoln | Mucolipin 2 | -2.12 | 0.023616 |
| Ccl5 | Chemokine (C-C motif) ligand 5 | -1.23 | 0.003339 |

Furthermore, when we analyzed enriched pathways in infected macrophages without HIF-α signaling compared to infected macrophages with HIF-α signaling, we found there were minimal significantly enriched pathways (Figure 4D). Together, this dictates there are HIF-α-dependent transcriptomic changes during homeostasis and in response to *L. major* infection supporting previous data depicting infection activates HIF-α. Finally, to further characterize the role of HIF-α signaling in infected macrophages under pro-inflammatory conditions, we compared the gene expression profile of infected macrophages deficient for HIF-α signaling stimulated with LPS/IFNγ to infected macrophages with intact HIF-α signaling under the same conditions. There were 102 DEGs between infected and stimulated macrophage without and with HIF-α signaling, 63 being upregulated and 39 downregulated (Figure 5A and Table 7). Of note, the top upregulated DEGs included *Slc26a11*, *Agap1*, and *Cxcr4* and the top downregulated DEGs contained *Mmgt1*, *Arhgap4*, and *Mdn1* (Figure 5A and Table 7). We predict the upregulated genes are inhibited by HIF-α signaling (*Slc26a11*, *Agap1*, and *Cxcr4)* while the downregulated DEGs are mediated by HIF-α signaling (*Mmgt1*, *Arhgap4*, and *Mdn1).* To identify pathways enriched for our DEGs, we performed a KEGG analysis. The KEGG pathway analysis revealed the DEGs clustered into pathways related to translation and protein production (‘Ribosomes’ and ‘Protein export’) (Figure 5B). Upregulation of the ‘ribosome’ and ‘protein export’ pathways in macrophages without HIF-α signaling suggests that these pathways are suppressed by HIF-α. To further conduct gene set enrichment analysis we utilized the molecular signature database (MSigDB). The MSigDB analysis revealed that the ‘interferon gamma response pathway’ was significantly upregulated in stimulated and infected macrophages without HIF-α signaling indicating this pathway is inhibited by HIF-α signaling (Figure 5C). Additionally, the ‘oxidative phosphorylation pathway’ was found to be upregulated indicating HIF-α signaling suppresses this pathway in infected macrophages in response to pro-inflammatory stimuli (Figure 5C).

**Figure 5:**
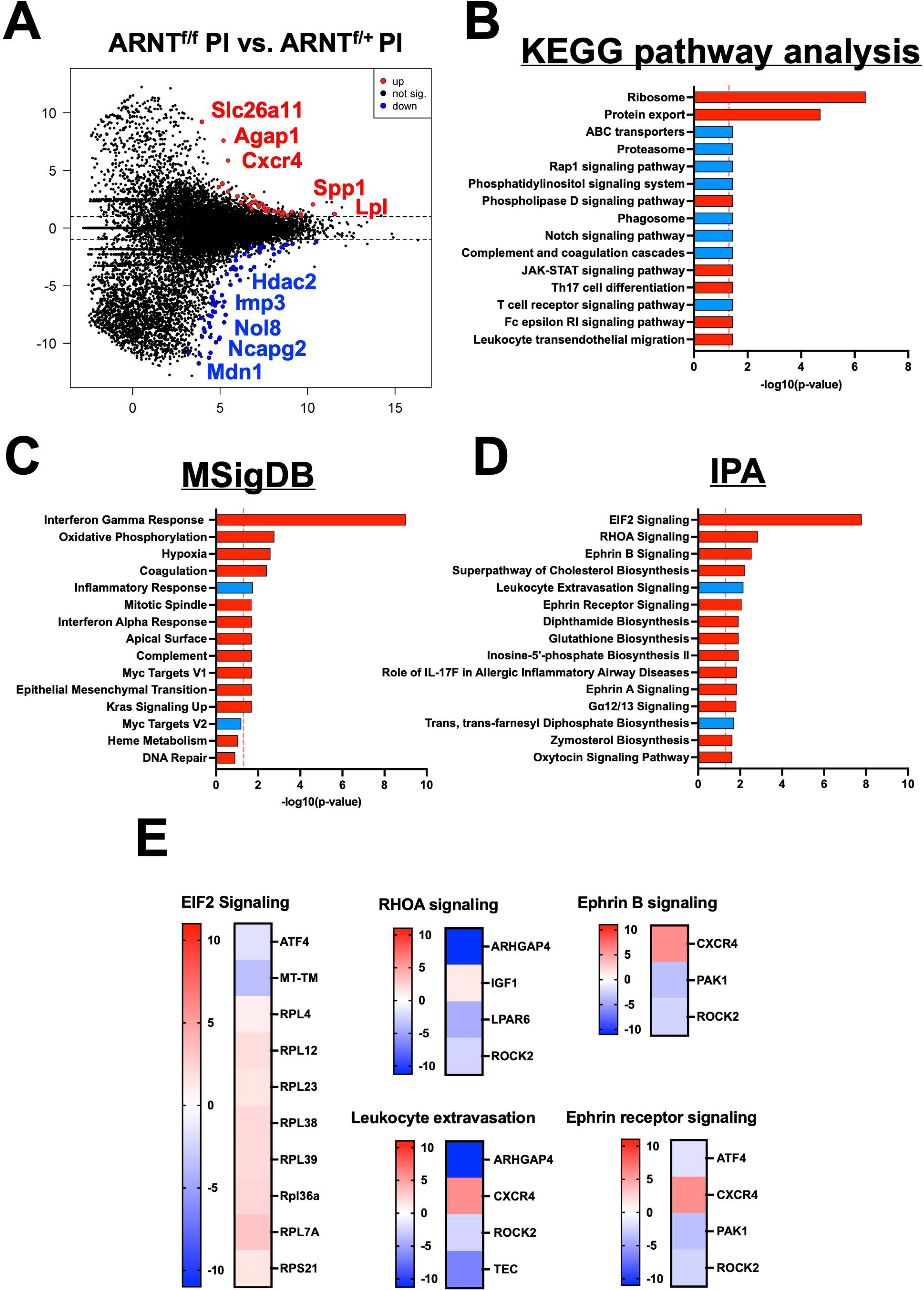
HIF-α signaling suppresses translational pathways under inflammatory conditions. RNASeq and pathway analysis on macrophages with and without intact HIF-α signaling infected with *L. major* and treated with pro-inflammatory stimuli. (**A**) DEGs upregulated (red) and downregulated (blue) in infected macrophages treated with LPS/IFNy without HIF-α signaling compared to macrophages with intact HIF-α signaling under the same conditions. (**B**) KEGG analysis identified enriched pathways in infected macrophages without HIF-α signaling stimulated with LPS/IFNy. (**C**) MSigDB pathway analysis defined upregulated pathways in red and downregulated pathways in blue in the infected macrophages without HIF-α signaling compared to macrophages with intact HIF-α signaling. (**D**) Ingenuity pathway analysis (IPA) was run to determine upregulated and downregulated pathways (red and blue respectively). (**E**) Heatmap plots of each upregulated or downregulated pathway defined by the IPA with individual altered DEGs represented in each pathway.

**Table 7.**
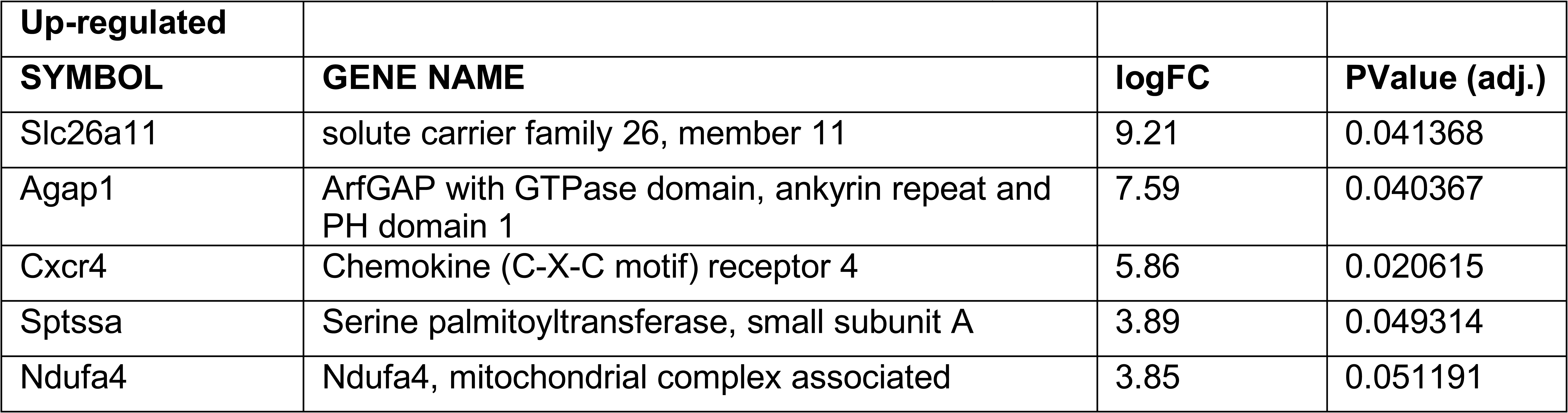
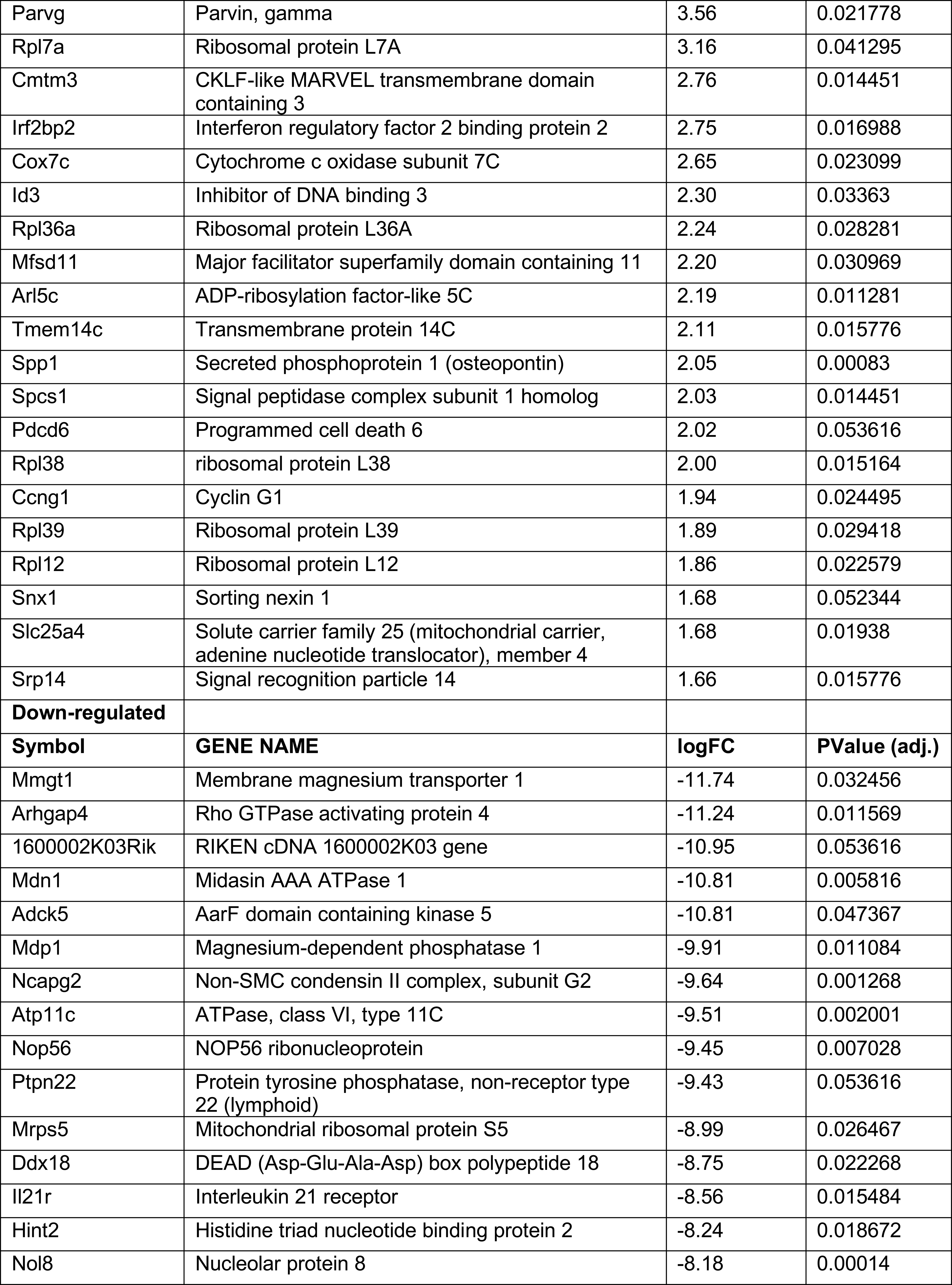
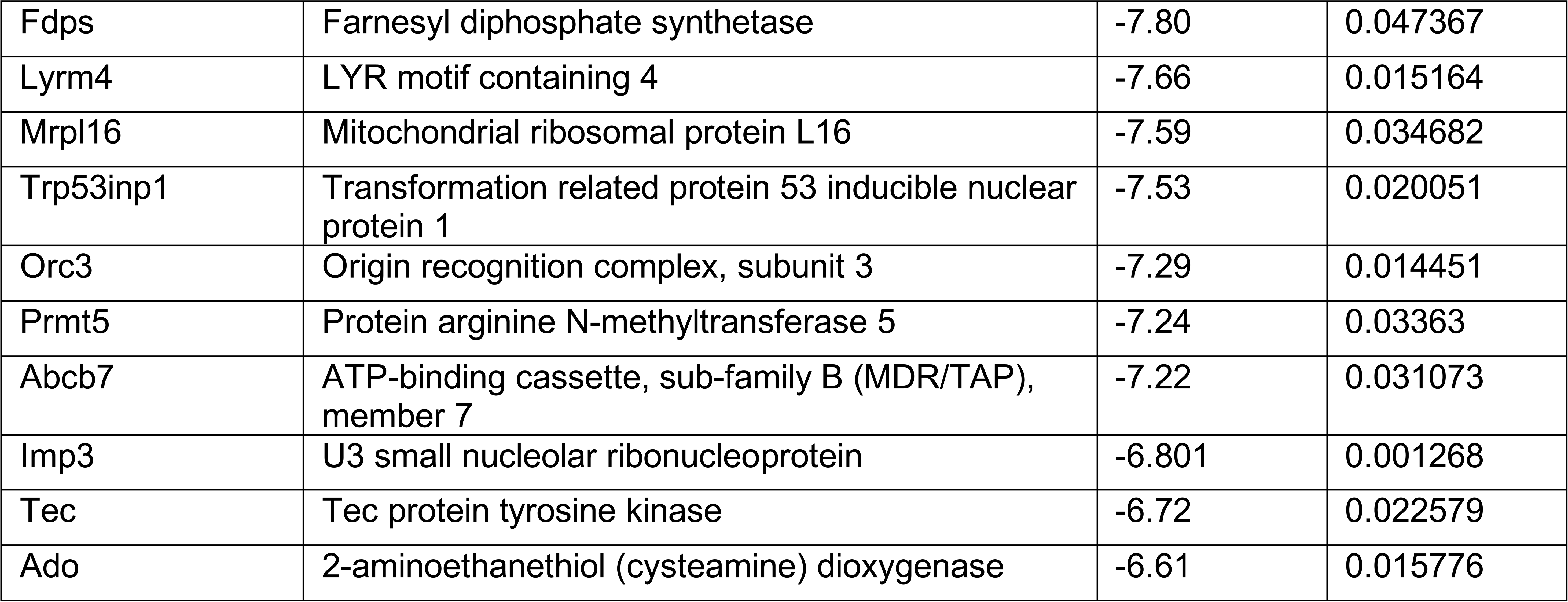
Top 25 Significantly Up- or Down-regulated DEGs between ARNT^f/f^ compared to ARNT^f/+^ Infected Macrophages Treated with LPS/IFNy.

Furthermore, many ribosomal related transcripts were enriched in the HIF-α deficient infected macrophages treated with pro-inflammatory stimuli such as *Rpl4*, *Rpl7a*, *Rpl12*, *Rpl23*, *Rpl38*, *Rpl39*, and *Rps21*. To further investigate these altered ribosomal transcripts, we conducted an ingenuity pathway analysis (IPA) comparing infected macrophages treated with LPS and IFNγ with or without intact HIF-α signaling (Figure 5D-E). We pinpointed ‘EIF2 signaling’ as the top upregulated pathway in macrophages without HIF-α signaling, proposing that in a scenario with both infection and inflammatory stimuli, HIF-α inhibits EIF2 signaling. This data is consistent with the KEGG pathway analysis indicating HIF-α signaling suppresses protein translation during inflammatory conditions. Of note, several other enriched pathways were identified by the IPA including ‘RhoA signaling’ and ‘Ephrin B signaling’ suggesting these pathways are inhibited by HIF-α (Figure 5D-E). Finally, ‘Ephrin Receptor Signaling’ and ‘Leukocyte Extravasation’ were downregulated in infected macrophages stimulated with LPS and IFNγ without HIF-α, again suggesting this pathway is mediated by HIF-α (Figure 5D-E).

## Discussion

HIF-α activation is a hallmark of both CL and VL occurring in response to tissue hypoxia, TLR activation, ROS and cytokines like TNFα and IL-1β, all of which are present during *Leishmania* infection (9,12,29,42–44). However, the direct contribution of the parasite versus the host response/microenvironment to HIF-α activation is not clear. *Leishmania* parasites can directly activate HIF-1α in macrophages, but the direct activation of macrophage HIF-1α is context dependent with the parasite species playing a major role. For instance, *L. amazonensis* parasites, which causes CL in South America, directly induces the expression of HIF-1α in human and mouse macrophages in vitro under normoxic conditions (10,45). HIF-1α is also present in *L. amazonensis*-infected skin (10). While *L. amazonensis* parasites can drive HIF-1α expression on their own, HIF-1α also promotes *L. amazonensis* killing by macrophages under hypoxic conditions (45). Similar to *L. amazonensis*, *L. donovani* parasites, which cause VL in Africa and Asia, directly activate HIF-1α in macrophages in vitro under normoxic conditions (46,47). *L. donovani* parasites increase HIF-1α expression, nuclear translocation and activity in a variety of macrophages including J774 cells, peritoneal macrophages and splenic-derived macrophages from BALB/c mice (46). To stabilize HIF-1α, *L. donovani* parasites use an array of mechanisms including depleting host iron pools to modulate prolyl hydroxylase activity and inducing microRNAs to limit NF-KB activation which establishes a suitable environment for parasite survival (46,47). In vitro, HIF-1α blockade inhibits *L. donovani* intracellular growth and HIF-1α stabilization promotes *L. donovani* growth inside macrophages (46,47). However, myeloid-specific HIF-1α^-/-^ mice infected with *L. donovani* and humans with a loss-of-function *HIF1A* gene polymorphism are more susceptible to infection (48). The role of HIF-2α in *L. amazonensis* and *L. donovani* infection has not been investigated.

Although *L. amazonensis* and *L. donovani* parasites can activate HIF-α directly, previous work shows that *L. major* parasites do not increase HIF-1α expression or activation under normoxic conditions in macrophages (11,23,27,29). For example, HIF-1α and HIF-2α as well as HIF-1α-specific and HIF-2α-specific target genes are increased at the site of murine *L. major* infection, but in vitro infection of macrophages with *L. major* does not induce HIF-1α expression (11,25). Rather *L. major* parasites require additional inflammatory signals such as LPS and/or IFNγ to induce HIF-1α accumulation and subsequent HIF-1α target expression like NOS2 and VEGF-A in macrophages (23,27,29). While HIF-α stabilization promotes *L. donovani* survival in macrophages, HIF-α stabilization does not impact *L. major* parasite growth in macrophages and may be why *L. major* parasites alone do not induce significant HIF-α protein accumulation (11,27,29). Here through pathway analysis, we have shown that in vitro, the HIF-1α signaling pathway is enriched during infection with *L. major* and many initial transcriptomic changes are HIF-α-dependent suggesting infection with *L. major* initiates the HIF-α transcriptional program (Figure 2). This is consistent with an additional study investigating initial transcriptomic changes after in vitro *L. major* infection, reporting HIF-1α signaling is enriched in murine macrophages at 4 hours post-infection (49). Despite these transcriptomic indications, it is possible pro-inflammatory stimuli are required for optimal HIF-α activation and subsequent target gene activation. It is important to note that in the above-mentioned study, *L. major* infection was not associated with changes specifically in HIF-1α accumulation. In the present study we have investigated changes in the absence of both HIF-1α and HIF-2α signaling which could account for the discrepancies.

HIF-α activation occurs in a wide variety of circumstances playing a central role in tissue adaptation to low oxygen tensions (50,51). Namely, hypoxia can be a characteristic of both tissue injury and subsequent inflammation, where infiltrating cells increase the demand for nutrients and oxygen, further depleting the tissue stores (52,53). Protein translation is an energetically demanding process and during hypoxia, inhibition of translation supports energy homeostasis and possibly promotes survival when energy stores are insufficient (54,55). Therefore, translation during hypoxic conditions becomes selective; coordinating adaptation to promote cell survival under low oxygen and energy conditions (56). Specifically, hypoxic conditions have been shown to stifle protein translation through downregulation of EIF2α signaling which we have shown is directly suppressed by HIF-α signaling in macrophages during inflammatory conditions (Figure 5) (57). Phosphorylation of eIF2a is necessary for mRNA translation inhibition during hypoxia and may be coordinated by HIF-α based on the current findings (55,58).

In addition to acclimating tissue to low oxygen availability, HIF-α is also a master regulator of macrophage inflammatory and innate immune function (15,59,60). Inhibition of protein translation coupled with a shift in metabolism to glycolysis during hypoxia are both mechanisms to conserve energy directly manipulated by HIF-α (61–63). Previous reports have demonstrated that HIF-α is capable of shunting macrophages towards a M1 dominant phenotype by targeting glucose metabolism (64,65). Elevated glucose metabolism coupled with HIF-α-induced ATP production are two major cellular mechanisms of overcoming low oxygen tension. As a result, macrophage-specific deletion of HIF-1α leads to impaired macrophage responses including lower glycolytic rates, lower energy generation, and impaired motility (66–68). Here we have shown that macrophages deficient for HIF-α signaling are predisposed to a dominant oxidative phosphorylation profile in comparison to HIF-α competent macrophages (Figure 5). Our study confirms that HIF-α reprograms macrophages during *L. major* infection to cope with the energetic demand. We have also shown through IPA analysis that pathways associated with macrophage motility are dysregulated during genetic deletion of HIF-α signaling including RhoA signaling and leukocyte extravasation consistent with what is reported in the literature (69–71).

In summary, we showed *L. major* infection elicits a subtle macrophage HIF-α program, but major transcriptomic changes dependent on HIF-α are only present in a pro-inflammatory environment. This supports our hypothesis that during in vivo *L. major* infection, HIF-α stabilization is dependent on the pro-inflammatory milieu and not *L. major* directly, which is the case during *L. donovani* infection (46). Additionally, we have evidence suggesting HIF-α suppresses protein translation in response to *L. major* infection and pro-inflammatory stimulus. Future work will investigate if protein translation is suppressed during in vivo infection with *L. major* in a HIF-α dependent manner and if this is unique to *L. major* or if it is conserved in other skin infections and diseases. We hypothesize suppression of translation is a mechanism of cellular adaptation to the pro-inflammatory response and subsequent hypoxic conditions from infiltrating cells and their high energetic demand during infection. A complete understanding of HIF-α during inflammation is vital in developing targeted therapeutics not only for CL but also for other inflammatory skin diseases as well as diseases where HIF-α is highly expressed.

## Acknowledgements

This work was supported by the Center for Microbial Pathogenesis and Host Inflammatory Responses (funded by NIH NIGMS Centers of Biomedical Research Excellence Grant P20-GM103625). This publication was also supported in part by funds provided by the National Center for Advancing Translational Sciences of the NIH under awards TL1 TR003109 and UL1 TR003107 for the Systems Pharmacology and Therapeutics (SPaT) NIH T32 training grant GM106999 to Lucy Fry. This study was supported by the Arkansas Children’s Research Institute, the Arkansas Biosciences Institute, and the Center for Translational Pediatric Research funded under the National Institutes of Health National Institute of General Medical Sciences (NIH/NIGMS) grant P20-GM121293 and the National Science Foundation Award No. OIA-1946391. The content is solely the responsibility of the authors and does not necessarily represent the official views of the NIH. The funders had no role in study design, data analysis, decision to publish or preparation of the manuscript.

